# Conformational landscape of full-length Smad proteins

**DOI:** 10.1101/2021.04.13.439655

**Authors:** Tiago Gomes, Pau Martin-Malpartida, Lidia Ruiz, Eric Aragón, Tiago N. Cordeiro, Maria J. Macias

## Abstract

Smad transcription factors, the main effectors of the TGFβ (transforming growth factor β) network, have been shaped along the evolution of multicellular animals to regulate essential processes. Smad proteins have a mixed architecture of globular domains and flexible linkers and adopt distinct quaternary structures depending on their activation state and cellular context. Here we studied the structures of full-length Smad4 and Smad2 proteins through an integrative approach combining small-angle X-ray scattering and detailed atomic information obtained from Nuclear Magnetic Resonance spectroscopy, X-ray and molecular dynamic simulations. Both Smad4 and Smad2 populate ensembles of expanded/compact conformations, with the MH1 and MH2 domains tethered by intrinsically disordered linkers that provide conformational freedom to the proteins. In solution, Smad4 is monomeric, whereas Smad2 coexists as monomer-dimer-trimer association states, even without activation. Smad2 dimers, which were previously overlooked, are proposed as key building blocks that define the functional quaternary structures of Smad proteins.

**Highlights:**

- Solution ensembles of full-length Smad2 and Smad4 proteins
- Smad4 populates dynamic extended and compact monomeric conformations
- Smad2 dimers may be the intermediate state to form heterotrimers with Smad4
- New integrative methodological strategy for examining multi-domain and flexible proteins

## Introduction

The TGFβ family of cytokines *(1)* regulates crucial cell specification processes that allow the large morphological diversity and adaptability of multicellular animals *(2, 3)*. The context-dependent role of TGFβ is modulated by the Smad (mothers against decapentaplegic) family of transcription factors, which transmit the TGFβ signals from the membrane TGFβ receptor to the nucleus (4, 5). Smad proteins are well conserved in all metazoans and are classified into three functional classes: Receptor-regulated Smad (R-Smad, 1/5/8 and 2/3), the Co-mediator Smad (Co-Smad or Smad4), and the inhibitors I-Smad (Smad6/7) *(6–9)*. Genetic inactivation of TGFβ receptors and Smad transcription factors facilitates tumor progression by reconfiguring the canonical TGFβ pathway in a disease-dependent manner *(10)*. This alteration triggers phenotypic plasticity like in the epithelial-to-mesenchymal transition, suppresses most anti-tumor roles of TGFβ signaling, and promotes metastatic colonization and progression *(11, 12).* Smad proteins, particularly Smad4, are highly mutated in tumors, with mutations spreading along their entire sequence, affecting the two well-structured domains (MH1 and MH2 domains) and also the unstructured linker (8). In cells, the basal state of Smad4 and Smad2 proteins is believed to adopt compact conformations, which could play an auto-inhibitory role, especially in tumor-mutated proteins *(13–15)*.

Modular proteins have been identified in all living organisms and they represent an evolutionary breakthrough that facilitated molecular diversity and function *(16)*. Many signaling proteins, such as Smad transcription factors, combine structured and disordered modules to enhance their functional versatility by defining specialized complex systems whose composition depends on the cellular context *(17–19)*. In fact, the function resides at both structured domains and linkers, which are not inert connectors. In the case of the Smad proteins, the linkers play active roles in Smad function as substrates for kinases and phosphatases and binding platforms for cofactors and ubiquitin ligases, which label Smads for degradation *(20–23)*.

However, the mixed architecture of globular domains and disordered-flexible linkers is a technical challenge from a structural perspective. Initial attempts to crystalize full-length Smad4 proved unsuccessful *(24)*, probably because the protein does not adopt a single globular structure. Thus, until now, all structural studies of Smad proteins have focused on the isolated domains and their complexes *(7, 25–32).* Although these fragments have revealed specific interactions in great detail, they only provide a partial picture of the function of full-length Smad proteins.

Studying the full-length structures of Smad proteins requires streamlined tools to unveil the ensemble of conformations that describe their dynamic behavior. Nuclear Magnetic Resonance (NMR), Small-angle X-ray scattering (SAXS) and molecular modeling provide truly complementary data, with detailed local conformations deriving from NMR data are merged with global shape/size fluctuations derived from SAXS. Combining these two methods in hybrid approaches that integrate data from high and low-resolution techniques is useful for analyzing the structure and conformational dynamics of these proteins and also compensates for the intrinsic limitations of each technique *(33)*. For instance, NMR is well suited to study dynamics and to map binding interfaces—even of large systems at residue resolution—but it is hampered by low sensitivity to increasing molecular weight. SAXS methods provide low-resolution information in solution but characterize dynamics and conformational equilibria independently of the protein size. In fact, similar hybrid approaches have recently been used for mixed flexible-rigid systems *(34–37)*.

We embraced the challenge of characterizing two representative full-length Smad proteins by combining SAXS with NMR data and high-resolution information available for their isolated domains. For the structural analyses of these proteins, we selected Smad4 (the common Smad), as it plays a pivotal role in both BMP- and TGFβ-activated pathways, and Smad2, as an example of a receptor-activated Smad of flexible protein assemblies in solution.

To analyze the SAXS data of Smad proteins, we built explicit models of the complete proteins, including monomers, dimers, and trimers. In our experimental conditions, the full-length Smad4 protein (Smad4FL) is mostly monomeric (and not a trimer as observed in crystals of its MH2 domain). The protein samples many conformations in solution, covering a range of inter-domain distances, with extended conformations being more abundant than compact ones. We also found that Smad2FL (with and without phosphomimetic mutations) associates as dimers and trimers through its MH2 domain. In the full-length context, the structured domains are flexibly linked, without retaining them through a long and stable compact interaction. Indeed, intrinsically-disordered linkers are known to play key roles modulating signaling proteins, but this dimension of Smad regulation has been missing until now due to the lack of structural information of the entire proteins.

The results presented here pave the way for the first description of the molecular landscape populated by full-length Smad proteins in solution and also for establishing how these conformations modulate the association with other Smad proteins and partners. Moreover, we believe that the strategy used in this work could be extended to investigate other transcription factors and modular signaling proteins that share with Smads the multi-domain architecture of globular domains separated by long flexible linkers and whose study has been hampered by technical difficulties.

## Results

### 1. General workflow

Prior to the analysis of the full-length protein datasets, we first studied the three main regions of the proteins. This information allowed us to build the pool of full-length models used to determine the conformational landscape of the proteins in solution that satisfy the experimental data. In brief, SAXS and NMR data were used to complete missing regions of the available MH2 domain structures and to characterize the linkers, both in isolation and in the full-length context. Based on the intrinsically disordered nature of the linkers confirmed by their NMR-fingerprint *(38, 39)*, we modeled them as flexible ensembles using the Flexible-Meccano pipeline *(40)*. We then selected sub-ensembles that collectively explain the SAXS data, applying the ensemble optimization method (EOM) *(41)*. Our previous description of the analysis of the MH1 domains in solution *(26–28)* supports the applicability of the complementary sampling strategy to describe flexible loops within well folded domains from SAXS and NMR data. Final models were selected using the EOM pipeline. Ensembles were represented using MH2 as reference (surface representation), with MH1 represented as spheres of variable diameter proportional to the frequency of a given inter-domain position within the ensemble. The general experimental workflow, including the constructs, are shown in Fig. 1A, B and fig. S1A,B.

**Fig. 1.**
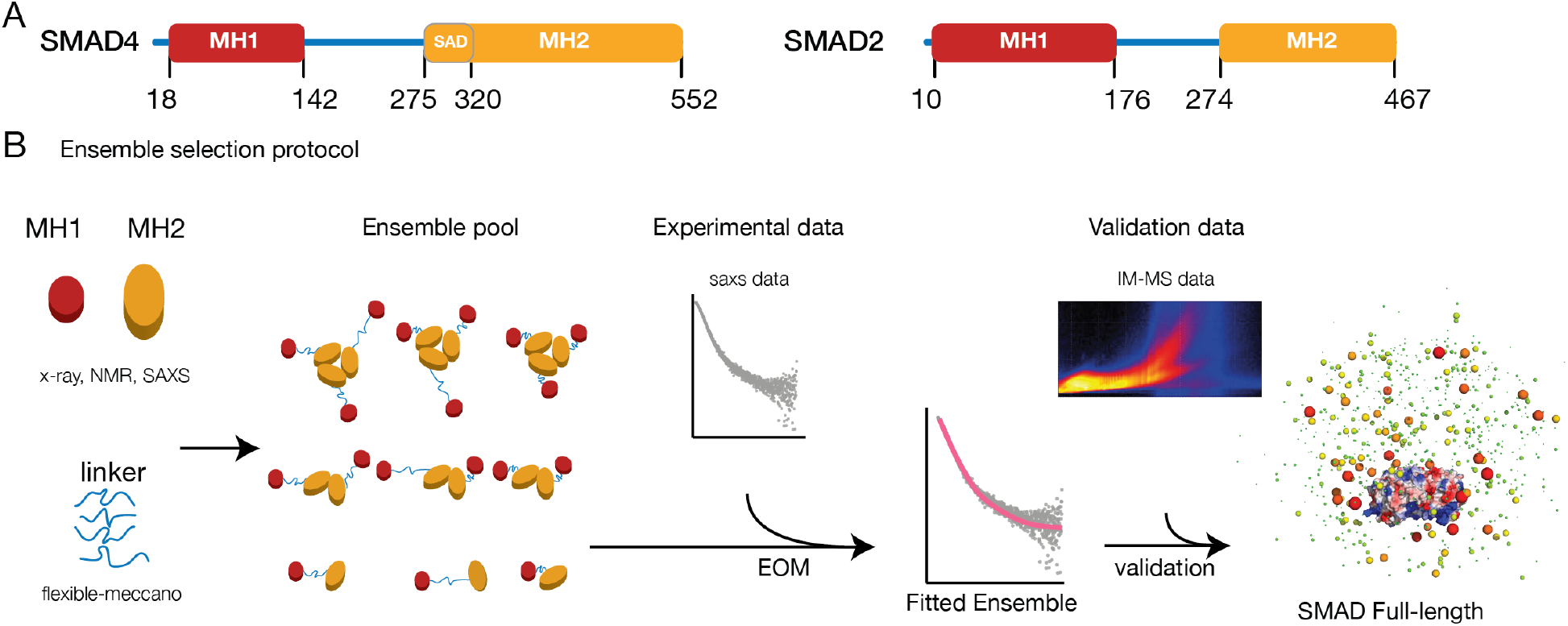
Construct design and general workflow **(A)** Smad4 and Smad2 domain composition. Detailed boundaries are shown as fig.S1A,B. **(B)** General methodology and the ensemble selection protocol. Models of the full-length proteins were prepared, including monomers, dimers and trimers. Linker extended conformations were generated using a *de novo* built ensemble. Flexibility of the linkers was verified by NMR. Final models were selected using the EOM pipeline. Smad4FL models were validated using IM-MS data. Final ensembles were represented using the MH2 as reference (surface representation), with the MH1 shown as spheres of variable diameter proportional to the frequency of a given inter-domain position within the ensemble.

### 2. R-Smad and Smad4 MH2 domains display distinctive assembly propensities in solution

The Smad4 and Smad2 MH2 domains form crystallographic trimers *(24, 42, 43).* Unfortunately, some regions in the domain were not defined in the electron density map due to flexibility. These regions involved the sequence connecting helices 3 and 4 and also the most C-terminal part of the domain. The addition of 46 residues preceding the Smad4 MH2 domain (named Smad Activation Domain, SAD) *(44)* showed that such an extension interacts with the domain core, stabilizing its structure in crystals, which also form crystallographic trimers. The SAD extension revealed the presence of a new strand (β9, residues 288–291 of the extended region, Fig. 1A, fig. S1A), a longer helix 3 than initially observed in the core domain, and two additional short strands at the C-terminus. However, about 50 residues located in loops were undetected in the SADMH2 domain, probably due to internal flexibility.

To analyze this dynamic behavior and to build a complete model of the domain, we purified three Smad4 MH2 constructs, the core MH2 domain (S4MH2) for NMR and SAXS studies, and two extended constructs, which were studied by SAXS. These constructs contain either the SAD region (S4SADMH2) or the full linker up to the MH1 domain (S4LMH2). The latter construct is similar to that previously studied for the Smad2 MH2 phosphomimetic domain in solution by SAXS *(45)* and it was built to study whether the presence of the linker modulates the association properties of the MH2 domain.

To build the MH2 domain models in solution, we analyzed the 3D backbone experiments of the S4MH2 domain acquired at 850 MHz using a fully deuterated and ^13^C- and ^15^N-labeled sample (residues 310-552). Since amide resonances corresponding to flexible areas are solvent-accessible, we re-suspended the fully deuterated protein in aqueous solution. We observed a rapid amide deuterium/hydrogen exchange and acquired 3D backbone experiments using non-uniform sampling. The backbone assignments confirmed that the helix 3 is flexible as long as it is observed in the S4SADMH2 structure, and that the loop connecting helix 3 and 4 involves residues 462-489 and does not adopt a secondary structure in solution. We also observed the presence of two short extended/β regions consistent with the boundaries defined for t1 and t2 in the S4SADMH2 structure (PDB:1DD1 *(42)* (fig. S2A). This information allowed us to use the structural elements of the extended MH2 domain as a template for all models, and building the remaining flexible regions as for the MH1 domains *(26–28)*.

The SAXS-derived distance distributions and Kratky plots corresponding to S4MH2 and S4SADMH2 constructs are characteristic of globular proteins, whereas the distribution of *Rg*-values of S4LMH2 (Fig. 2A, fig. S2A) corresponds to a broader family of conformers than expected for a folded MH2 domain tethered to a pure random coil linker. Regarding the analysis of the data, the starting pool of models contains monomers, dimers and trimers (10,000 conformers of each class). The ensemble optimization method (EOM) was applied to select those models that fit the experimental SAXS data at different concentrations. Independently of the linker length and concentrations, the selected models that fit the SAXS data well contain only monomeric species (S4MH2: χ^2^=0.77, *Rg*=22Å, S4SADMH2: χ^2^=0.87, *Rg*=25Å, S4LMH2: χ^2^=0.63, *Rg*=37Å) (Fig. 2A). In fact, when the pool contained only trimers (as in crystals), the generated curves did not fit the experimental data (χ^2^=21, fig. S2B). Furthermore, the presence or absence of the SAD domain or of the linker preceding the MH2 domain did not affect the quaternary structure of the fragments at different concentrations, which were always monomeric (fig. S2B-C).

**Fig. 2.**
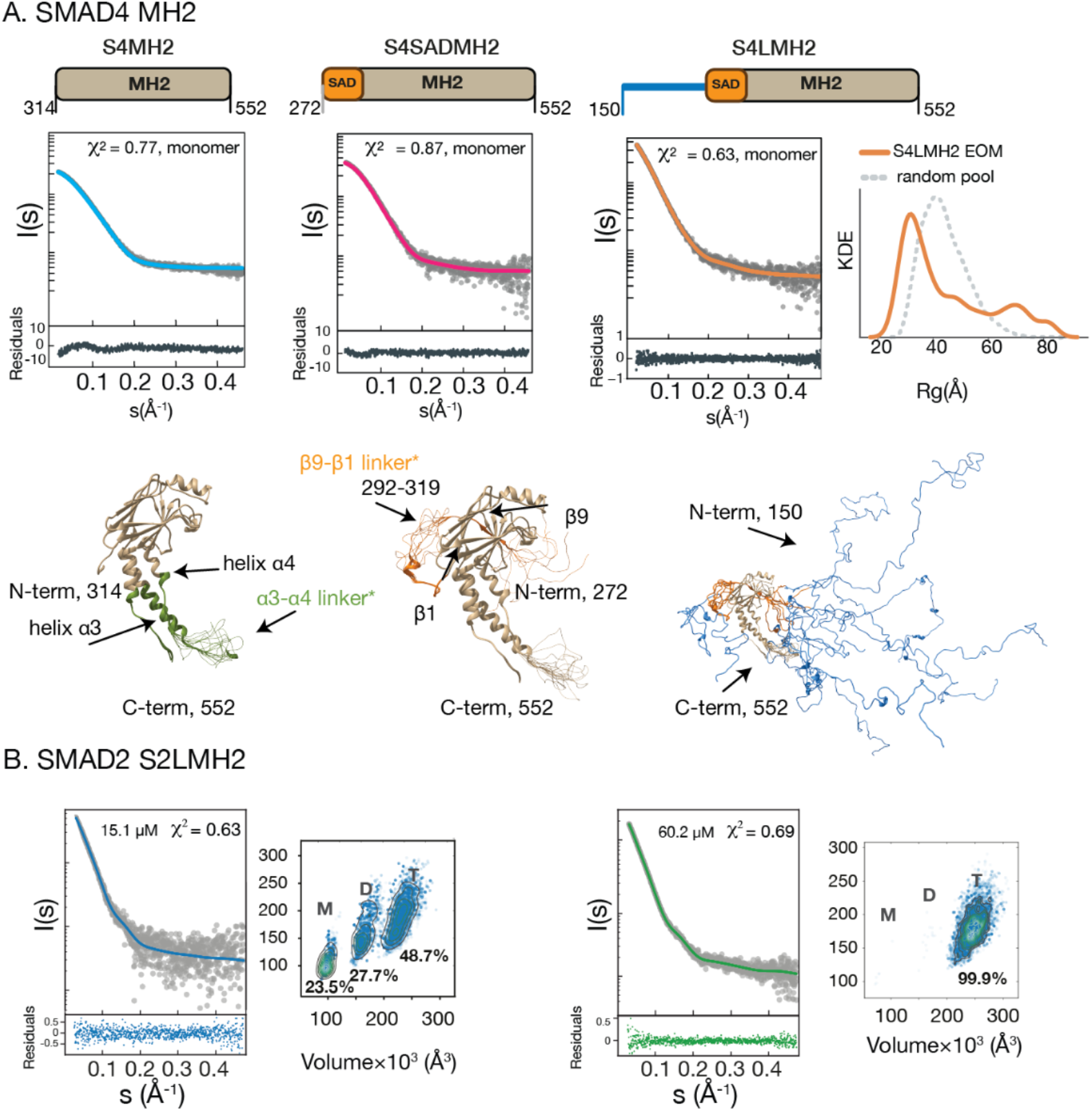
Smad4 and Smad2 MH2 domains in solution **(A)** Three different MH2 domain constructs of Smad4 proteins studied using SAXS. SAXS intensities I(s) (in gray) are represented with respect to the momentum vector s for the simulated profiles. The S4LMH2 SAXS curve was acquired at 29.7 μM, the SAXS scattering curves for S4MH2 and S4SADMH2 correspond to a merged curve generated from data at several protein concentrations (S4MH2:19-790 μM; S4SADMH2: 32–540 μM). Fittings are shown in blue, red-pink and orange, for each MH2 domain construct. Residuals showing the agreement between the simulated and experimental profiles are given below each curve. In the case of the S4LMH2 construct, which contains the flexible linker, we provide the Kernel Density Estimation (KDE) calculated for each EOM model (orange curve) superimposed to that obtained using the random pool of models (gray). Explicit models satisfying these curves are shown below the SAXS curves. Green regions in the S4MH2 ensemble depict NMR-supported secondary structures, which were not observed in X-ray structures. Orange regions in the S4SADMH2 are modeled based on secondary structure propensities and predictions. All selected models are monomeric. **(B)** SAXS data corresponding to the S2LMH2 non-phosphorylated domain at two concentrations. In these cases, monomers, dimers and trimers (at low concentration) or mostly trimers (high concentration) contribute to the ensemble of conformations in solution.

With respect to the Smad2 MH2 domain, we focused our analysis on the extended construct of the non-phosphorylated Smad2 MH2 domain (S2LMH2, Fig. 2B), which has not been previously studied in solution. The EOM-ensemble analysis suggests that this Smad2 domain coexists as a monomer-dimer-trimer equilibrium, with 50% of the total population organized as a trimer already at concentrations of 15 and 30 μM, and with the trimer being the main species at 60 μM (χ^2^=0.69).

Overall, our data reveal that besides the sequence and fold conservation between the Smad4 and Smad2 MH2 domains (39% and 51% identity and similarity, respectively), their MH2 domains display distinctive assembly propensities in solution. Whereas Smad4 is monomeric, Smad2 populates a pre-existing trimeric state even in the absence of a phosphorylation-dependent trigger.

### 3. The linkers behave as intrinsically disordered regions

The inter-domain sequences (of approximately 100-120 residues) are characteristic of each type of Smad protein *(8, 46)*. There is no structural information available for these linkers (Linker Segment, LS), except for short fragments of Smad1 and Smad2 in solution, which were studied as complexes bound to WW domains *(23)*. The analysis of the sequences of these proteins indicates that both LSs have a flexibility propensity, net charge and hydrophobicity propensities similar to those often defined as Intrinsically Disordered Regions (IDRs) *(47)* (fig S3A,B).

To decipher the dynamic properties of these regions, we started by analyzing the backbone and ^15^N T1, T2, and heteronuclear Overhauser effect (hetNOE) NMR relaxation experiments using recombinant samples. The 2D Heteronuclear single quantum coherence (HSQC) NMR spectra showed most of the amide signals clustering between 7.5 and 8.6 ppm but the signal overlap is not severe, allowing us to identify most residues in the linkers (fig. S3C). Chemical shifts confirmed the lack of secondary or tertiary structure. The T1 and T2, as well as the low hetNOE values (below 0.7) and the absence of secondary structure propensities, are all in agreement with values reported for flexible regions (Fig. 3A), *(48–50)*. The N-terminal part of S2L, which is adjacent to the MH1 domain, has a β-sheet/extended propensity, perhaps due to the high Pro content of this sequence, albeit of a flexible nature as indicated by the hetNOE values below 0.2. We also acquired a ^1^H-^15^N HSQC spectrum for Smad4FL and superimposed it to that of S4LS (fig. S3D). Over 75% of the Smad4FL visible resonances overlapped between the two constructs, with low dispersion for the ^1^H dimension, thereby indicating that the linker maintained a similar IDP-like behavior when isolated and in the full-length protein context. Quantitatively, the per-residue distribution of the hetNOE and T2 relaxation values for S4FL resemble those of S4LS (Fig. 3A), thus reinforcing the flexible character of the linker within S4FL. Unfortunately, Smad2FL could not be studied due to protein oligomerization and precipitation at the concentrations required for NMR.

**Fig. 3.**
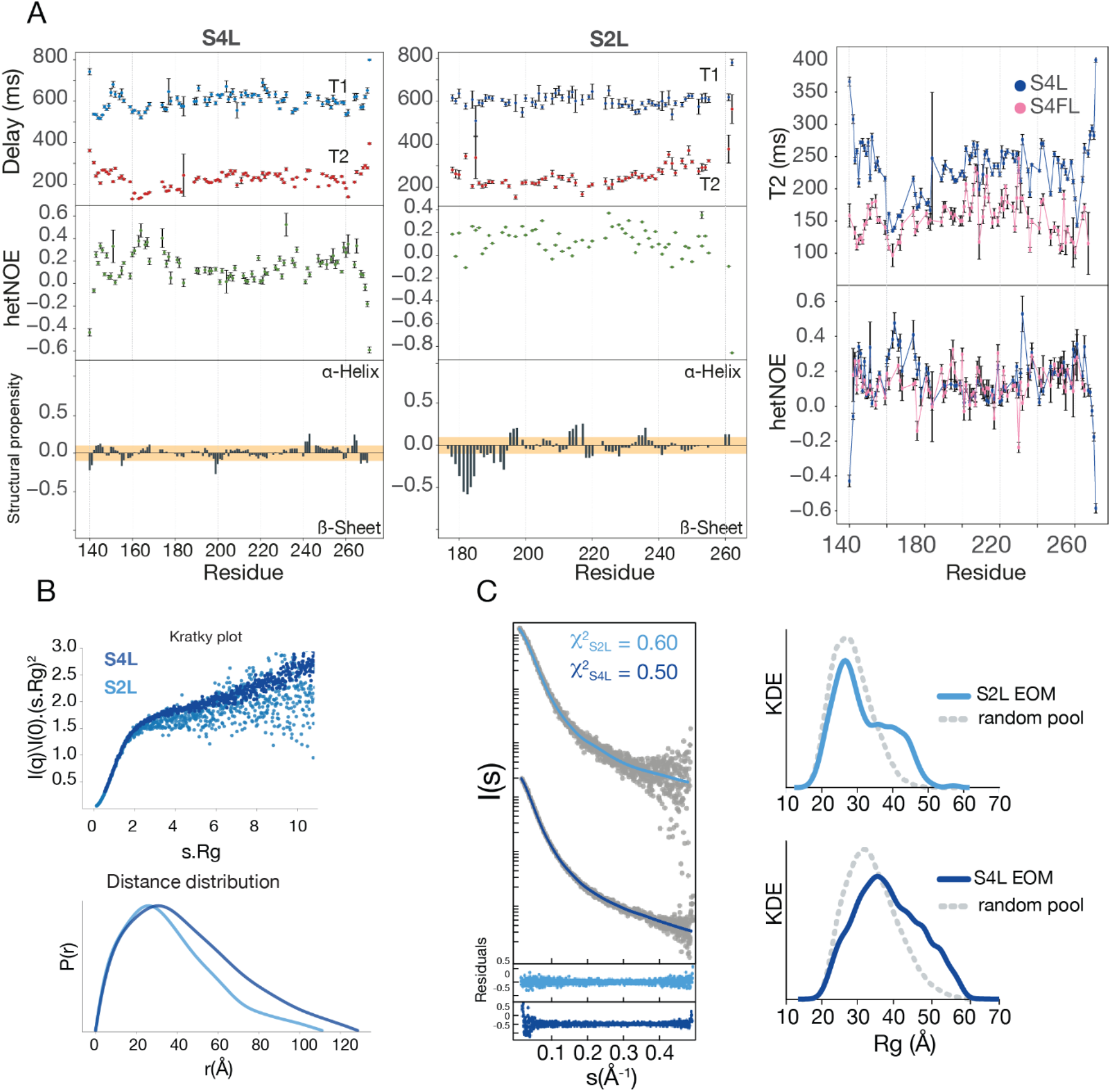
SAXS and NMR data of inter-domain linkers in solution **(A)** S2L and S4L relaxation experiments and secondary structure propensities. The comparison of the spin-spin relaxation time, T2, and the hetNOE for S4FL and S4L. Structural propensities were calculated using ncSPC *(50)*. The yellow bar depicts the random-coil threshold and values above or below the bar represent secondary structure propensities (ɑ-helix or β-sheet/extended, respectively). For IDPs, values close to β-sheet propensity imply that these IDPs have an elongated structure. The ^1^H-^15^N HSQC spectra showing the narrow 1H chemical shift dispersion characteristic of IDPs is shown as fig. S3B. **(B)** Kratky plots for S4L and S2L are shown in dark and light blue respectively. The high flexibility of both linkers is observed as a monotonic increase along with high s values. Distance distributions derived from the SAXS experimental profiles for S2L and S4L. The color code is the same as for the previous panel. **(C)** The experimental SAXS profiles and KDE for *Rg* distribution for S4L and S2L (in gray) and the solid line simulated curves from each EOM with residuals represented below.

We also monitored the degree of linker flexibility using the SAXS derived Kratky plots, which are consistent with an IDR profile (Fig. 3B). Each Kratky plot monotonically increases without a clear maximum, indicating conformational heterogeneity and lack of globularity *(51)*. The asymmetric pair distance distribution functions, *P(r)*, obtained from their scattering data, are also compatible with unfolded particles in solution, having a radius of gyration (*Rg*) of 29.9±0.4 Å and 36.8±0.2 Å, for S2L and S4L, respectively. These *Rg* values are close to theoretical ones for fully disordered random coils (*Rg*_S2L_^Rc^ = 27 Å, *Rg*_S4L_^Rc^ =33 Å) and they are identical for a polymer random coil of equal sequence length (fig. S3B). Their *P(r)* functions smoothly end at maximum distances (*Dmax*) of 111.0±2 Å for S2LS and 128.5±2 Å, for S4LS.

To further characterize the conformational properties of the linkers, we built large ensembles (10,000 conformers) for each isolated linker sequence, using the *Flexible-Meccano* (FM) algorithm *(40)*. Ensembles of this size are recommended for highly flexible systems like IDRs *(52)*. For each pool of conformations, we ran the EOM to yield sub-ensembles that reproduce the scattering profiles. The resulting subsets display *Rg* distributions enriched in conformations with slightly larger *Rg* values than a theoretical random coil, fitting the experimental SAXS curves *(53)* with χ^2^ values of 0.6 and 0.5 for S2L and S4L, respectively, and with better residual distribution than that corresponding to theoretical random coils (Fig. 3C). We observed that these dynamic properties were unaffected by the presence of the structured MH1 and MH2 domains in the full-length protein. By simultaneously fitting a single S4FL EOM pool (truncated as required for each SAXS experimental profile) to data acquired on the equivalent experimental constructs, we confirmed that linker conformations have similar overall flexibility profiles when in isolation, attached to the MH2 domain, and in the context of the full-length protein, adopting slightly more compact conformations than those expected for a theoretical random coil distribution. This multi-curve fitting approach yielded excellent χ^2^ statistics of 0.78, 0.95, and 0.76 for S4L, S4LMH2, and S4FL, respectively (fig. S3E).

Taken together, these results showed that Smad linkers behave as IDRs and that, in the case of Smad4, this behavior is observed in both isolation and in the full-length protein context as indicated by NMR and SAXS data.

### 4. Full-length Smad4 is a monomer and populates multiple conformational states in solution

To gain insights into the conformational landscape of full-length Smad proteins, we acquired SAXS data for Smad4 and Smad2 at different protein concentrations (Fig. 4A).

**Fig. 4.**
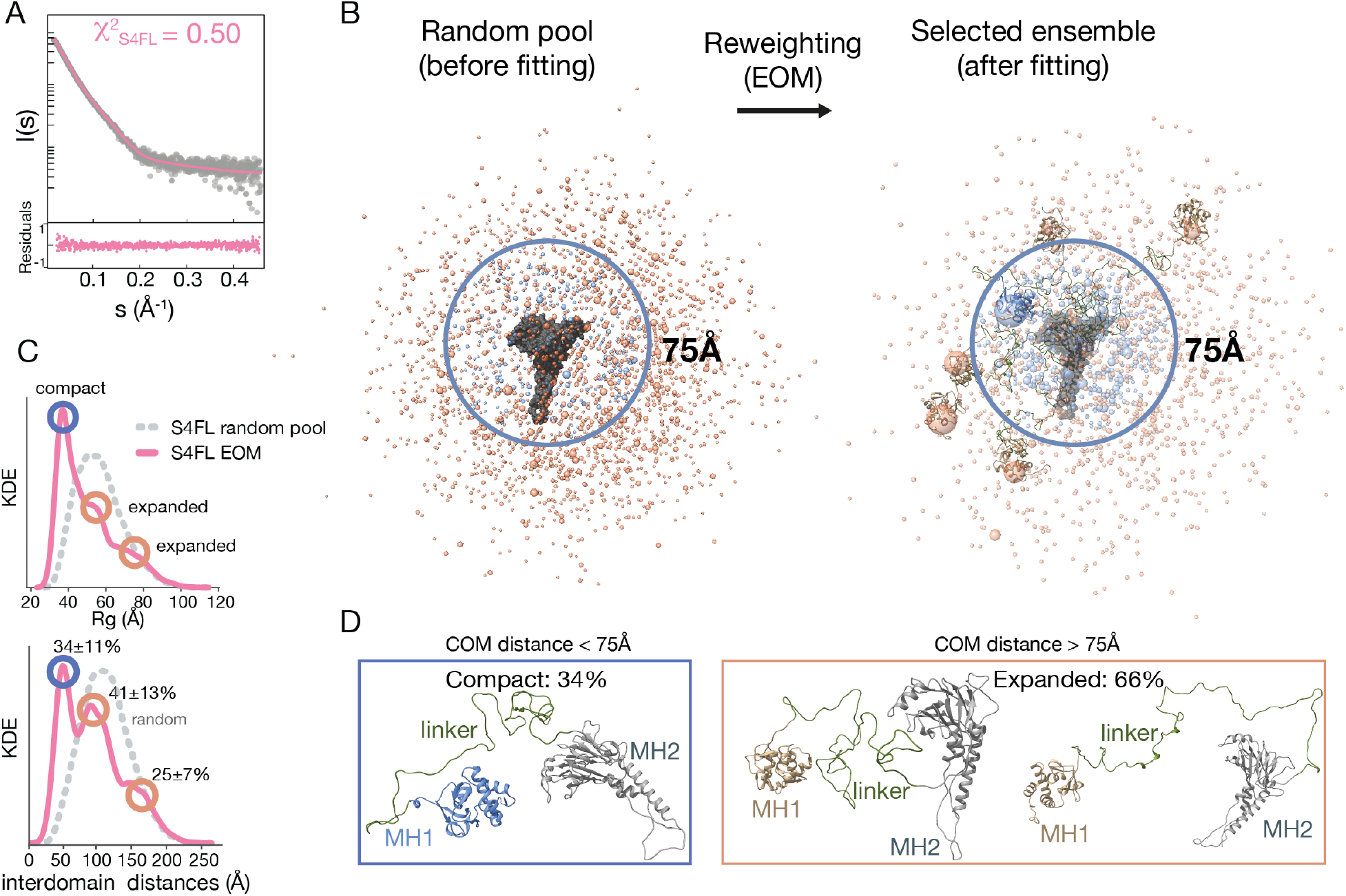
S4FL conformational landscape in solution **(A)** The SAXS EOM simulated profile corresponding to S4FL is shown in pink, and it is overlaid with the experimental profile in gray and the respective residuals at the bottom panel. **(B)** EOM and random pool S4FL conformational landscapes. To facilitate the identification of the domains, the MH2 domain is depicted as a surface (gray) whereas the MH1 domain is simplified as a sphere whose radius is proportional to the probability of occurrence for a given conformation. Blue spheres represent distributions up to a distance of 75Å between domains, and spheres colored in tan indicate expanded conformations. **(C)** Size distribution of the ensemble of conformations in the random pool and after EOM selection. The distribution shows compact (34%) and expanded (66 %) conformations, respectively. **(D)** Most representative conformations are indicated as explicit models.

In solution, S4FL shows a moderate flexibility as reflected by the asymmetry of the SAXS-derived P(r) distribution (*Dmax*=171.4Å and *Rg*=47Å) intermediate values between IDR and globular structures, as expected from the mixed architecture of the protein (fig. S3B). The Kratky plot also shows a plateau characteristic of modular proteins separated by flexible linkers *(54)* (fig. S4A).

To describe the ensemble of conformers that satisfy the experimental SAXS restraints, we generated explicit models built using structured MH1 and MH2 domains tethered by a linker modeled as an IDR. The models were built to include monomer, dimer and trimer populations to generate an ensemble pool containing 10,000 models with inter-domain distances covering a large range of spatial distributions as experimentally characterized. Sub-ensembles that collectively fit the SAXS curve were selected using EOM. The minimal sub-ensemble size (N_se_) of 50 models was empirically determined by searching for the smallest size where increasing the N_se_ did not show a significant improvement in χ^2^, to avoid overfitting (fig. S4B). Analysis of these ensembles revealed that, as with the S4MH2 constructs, only monomers were compatible with the experimental data.

Since the linker can adopt a variety of conformations, we grouped the EOM-selected models on the basis of the inter-domain center-of-mass (COM) distances, an approach that resulted in three major clusters with distances of approximately 50Å, 100Å, and 160Å (Fig. 4B,C). These clusters correspond to *Rg* values of around 35Å, 50Å and 70Å, respectively, resulting in a compact to expanded ~1:2 ratio. When compared to the random ensemble, which followed a Gaussian distribution centered at inter-domain distances of about 115 Å and *Rg* of ~55 Å, these experimental clusters are more compact (Fig. 4C,D). It should be noted that the three clusters that satisfied the experimental SAXS data display inter-domain distances too far to allow for direct MH1 and MH2 interaction, even for the most compact one, which has an inter-domain COM distance of 50Å. The other two clusters show very large values with an average separation of approx. 93 and 155Å. The compact/expanded ratio was validated by IM-MS, where monomers and similar ratios were observed for various m/z (fig. S4C).

To visualize the ensemble of models (pool and EOM-selected), we overlaid them with respect to the MH2 domain (surface representation). MH1 domains are represented as spheres condensed at the COM and the linkers are omitted for simplicity. In the pool, all spheres representing the MH1 domain have a similar diameter and are uniformly distributed. In contrast, in the ensembles that fit the experimental data, the spheres have different diameters. More populated regions are represented by larger spheres (Fig. 4B) since volumes are proportional to how often a given domain populates one region in the conformational space around the MH2 domain. Regions with an inter-domain distance of up to 75Å are shown in blue and the rest in tan.

The final ensemble agreed with the experimental data for models corresponding to monomeric proteins with a χ^2^=0.50 and a random dispersion of the residuals. Some representative models of compact and extended conformations are indicated (Fig. 4D). As expected, the pool ensembles are unable to explain the experimental SAXS data (χ^2^=1.98) (fig. S4D).

We used NMR titrations to further explore the potential interaction between the MH1 and MH2 domains in solution using isolated MH1 and MH2 domains (up to 3 equivalents, fig. S4E). The analysis did not reveal significant differences in the resonances (or intensity changes) in the MH1 domain, thereby indicating that the MH1 and MH2 domains do not form stable complexes in solution in the context of the full-length protein or of isolated domains.

These results indicate that, in solution, S4FL is a monomer and it populates an ensemble of conformations following a compact to expanded 1:2 ratio (Fig. 4A,B).

### 5. Smad2 exists as a monomer-dimer-trimer equilibrium, shaped by phosphomimetic mutations

To explore the conformational equilibrium of Smad2 in solution and its dependence on MH2 domain activation, we acquired SAXS data on both Smad2FL (S2FLWT) and also on a phosphomimetic Smad2 (S2FLEEE) described in the literature *(55)*. Data were acquired at different concentrations, (13.6, 24.3, 46.8, and 56.1 μM, for S2FLWT and 9.4, 18.7, and 28.1 μM for S2FLEEE, Fig. 5A,B, and fig. S5A,B), and analyzed using explicit ensemble models including monomer, dimer and trimer populations based on MH2 domain contacts. Dimers have been described previously *(56)* and our molecular dynamics simulations also indicated that the dimer is stable (fig. S5C). All χ^2^ values for the EOM-selected ensembles sustain the agreement with the SAXS data for both variants. In the case of S2FLWT, the trimeric state ranges from 1.0% at 13.8 μM to 16.4% at 56.1 μM, whereas for the S2FLEEE variant, the trimer population is 44.8% at 28.1μM. In both Smad proteins (Fig. 6 and Fig. 7), trimers are enriched at higher concentrations and their formation is enhanced by the phosphomimetic mutations. The EOM-selected models are shown using similar representations as for S4FL.

**Fig. 5.**
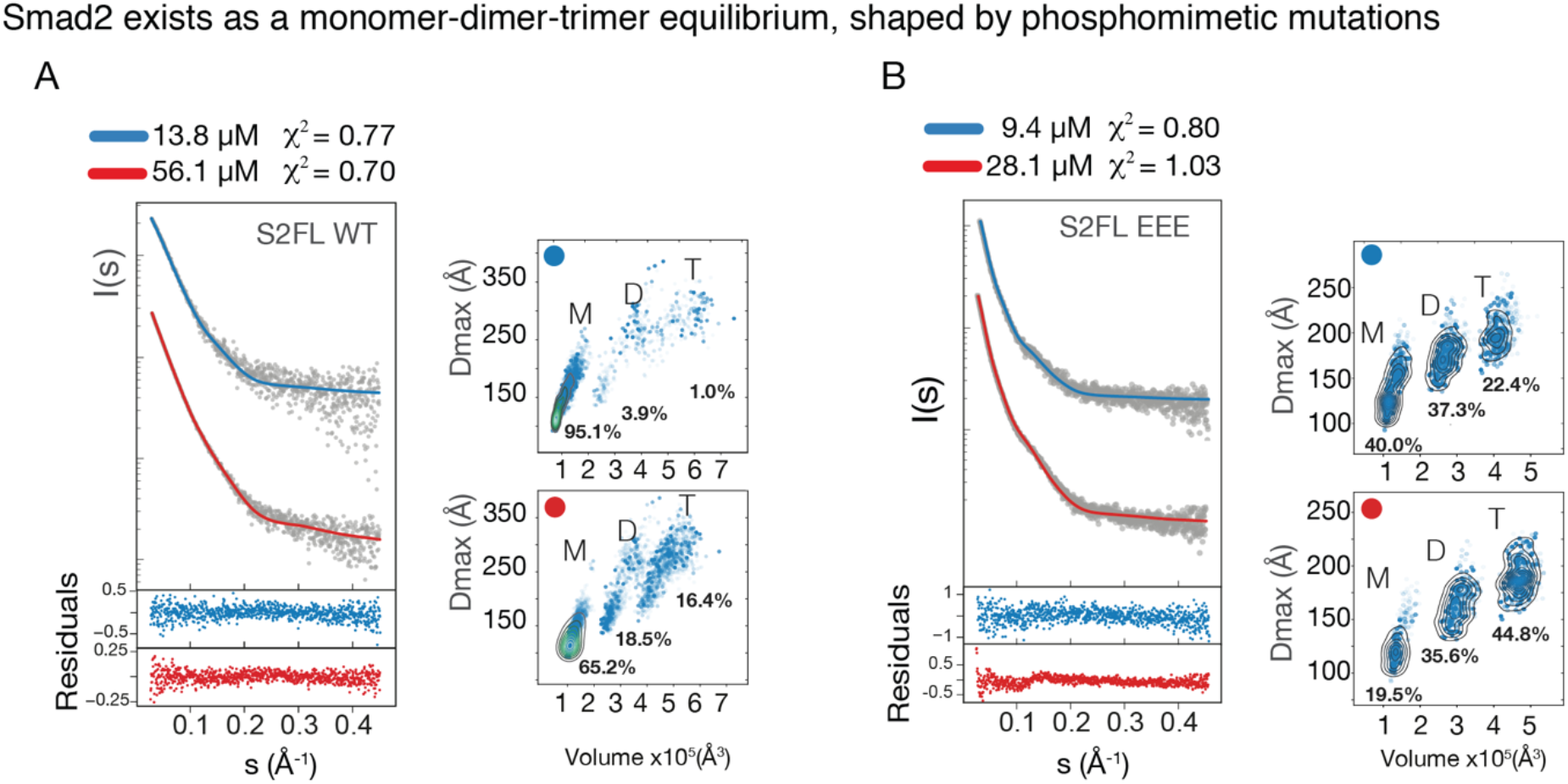
S2FL equilibrium distribution in solution **(A** and **B)** SAXS curves for S2FL WT or S2FL EEE at two different concentrations in gray and EOM fittings in blue and red. Next to each SAXS curve are the Kernel density contour plots for *Dmax* and Volume, calculated from the EOM ensembles. M, D and T are abbreviations for monomer, dimer and trimer species, respectively.

**Fig. 6.**
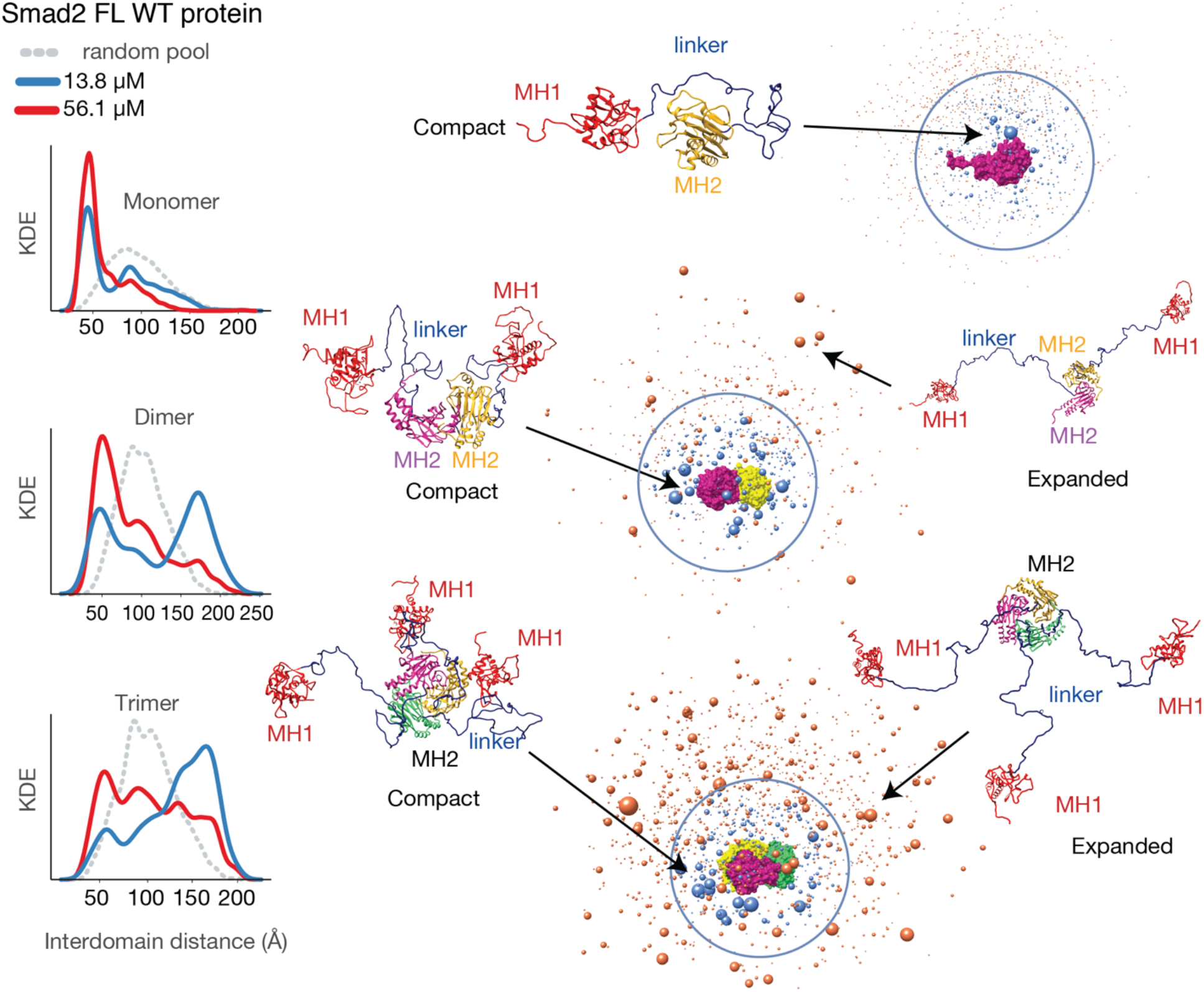
S2FL WT conformational landscape in solution. **Left**: MH1-MH2 inter-domain distance distribution of the pool ensembles compared to those obtained after the EOM-selection corresponding to the S2FL WT protein. Compact structures were classified as those with inter-domain distances < 75 Å. **Right**: Models derived from SAXS data at 56.1 μM are shown following a similar approximation as that used for S4FL proteins. Representative conformations (indicated with arrows) are depicted as explicit models. To facilitate the identification of monomers, dimers and trimers present in S2FL, the MH2 domains have been colored in purple, green, and yellow, whereas the MH1 domains are shown in red.

**Fig. 7.**
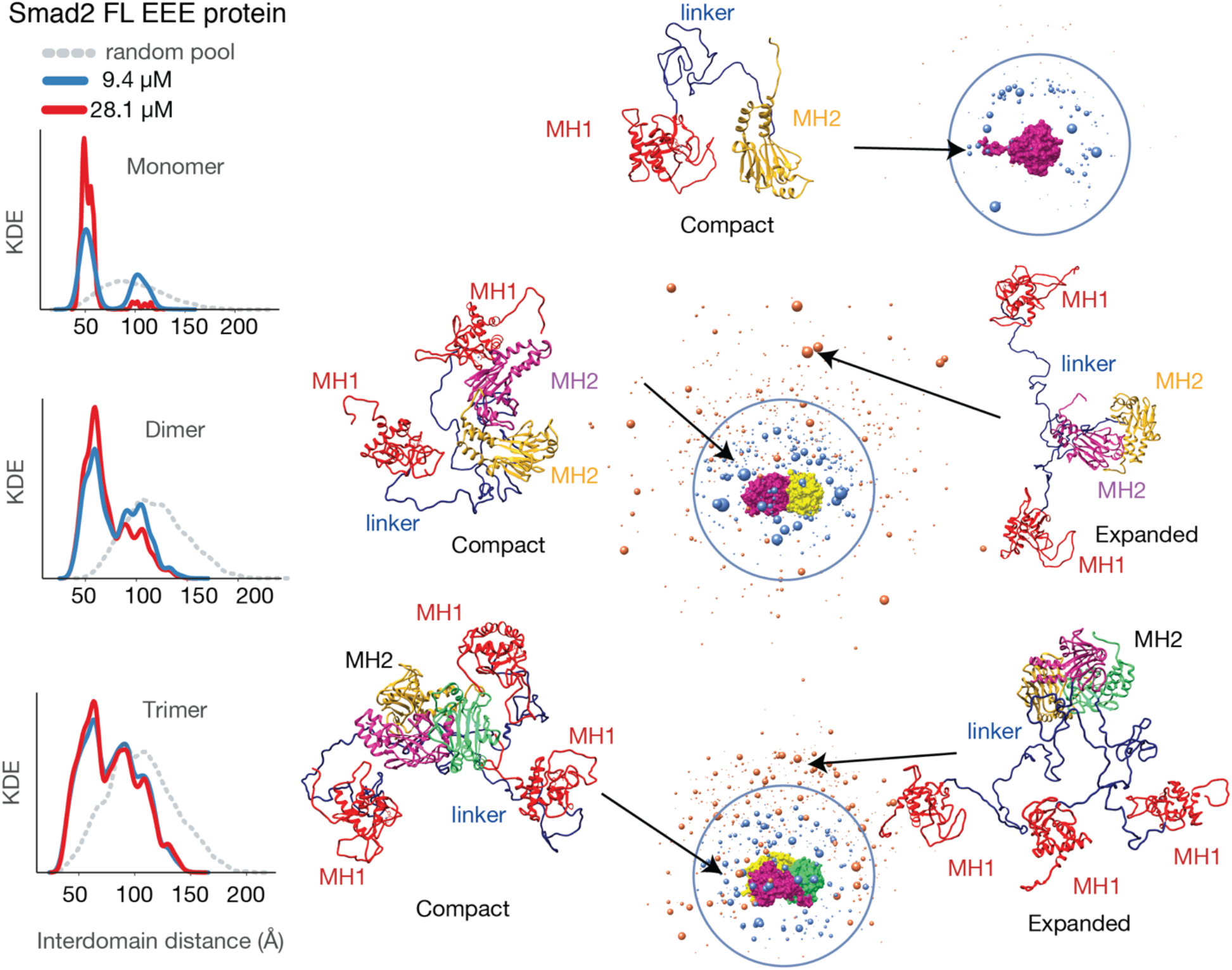
S2FL phosphomimetic variant conformational landscape in solution Inter-domain distance distribution and models derived from SAXS data corresponding to S2FL EEE protein. The representations are prepared following the same representations as in Fig. 6 and derived from SAXS data at 28.1 μM.

Remarkably, the dimer formation reaches a maximum of 18% at 56 μM in S2FLWT while for S2FLEEE dimer populations are almost invariant with concentration, with a constant fraction of approximately 36% at all the concentrations analyzed (Fig. 6 and Fig. 7). This concentration independence suggests that dimer formation is an intermediate in the monomer-trimer equilibrium. The presence of dimers might be crucial to ensure the formation of heterotrimers with Smad4 proteins in native conditions.

As a control of protein association, we prepared a mutant that contains a premature stop codon at position 460*, which produces a protein without the phosphorylatable region (S2FL_460*_ Fig. 1A). This variant is listed in the COSMIC database as being present in some tumors *(57)*. For S2FL_460_, almost no trimer (4%) was observed at high concentrations (74 μM), thus confirming the essential role of the C-terminal residues for trimer formation (fig. S5D). Abolishing trimer formation and consequently TGFβ activation could indicate that tumors harboring this change inactivate TGFβ and its tumor-suppressor phenotype *(7, 58)*.

We also explored the inter-domain distances, similarly to what we did for S4FL, using the EOM-derived ensembles. We found that an increase in protein concentration shifted these distances towards more compact arrangements, especially in S2FLWT (Fig.6 and S5E). Remarkably, the ≈50Å inter-domain distance resembles that of S4FL (Fig. 4A-C), despite the different inter-domain linker length (~40 amino acids shorter than in Smad4).

## Discussion

Proteins and domains are often represented as high-resolution static models. However, biomolecules in native contexts adopt dynamic ensembles of conformations, covering structural fluctuations from minor local movements to large-scale changes. The dynamic behavior of proteins, particularly of modular proteins like Smads, is crucial for their activity in the context of living organisms because flexibility provides an additional switch to fine-tune their structures to specific functional needs. In the case of Smad proteins, local flexibility at the global domains have been observed at the DNA binding interface of the MH1 domains and in loops and helices of the MH2 domains. Local flexibility in the linkers facilitates prolyl *cis*-*trans* isomerization, phosphorylation and also exposing primary contact sites to binding to activators and ubiquitin ligases for degradation *(23, 26, 27, 59)*. Studying global structured/disordered interplay has proved challenging in Smads due to the long linker that connects the two well-structured domains. We tackled this challenge through an integrated structural biology approach while studying the conformational landscape and self-assembly properties of Smad4 and Smad2 proteins. We observed that in addition to local flexibility at the domain level, Smad proteins also undergo large conformational changes that affect their overall shape distribution. In the absence of cofactors, the MH1 and MH2 domains are flexibly linked, allowing the domains to sample different relative orientations with respect to one another, without populating compact and globular architectures. The linker appears to act like a mechanical element, allowing the approximation and separation of the MH1 and MH2 domains without retaining them through a long and stable compact interaction. These transitions between compact and open states are usually favored for many multi-domain DNA-binding proteins *(37, 60, 61)*, and in the case of Smad proteins, the expanded states could facilitate linker modification and cofactor association by providing ensembles of conformations separated by almost no energy barriers.

Moreover, we observed that Smad4 is mostly monomeric (and not a trimer, as observed in crystals of MH2 domains) whereas Smad2 populates an equilibrium containing monomers, dimers and trimers, with the trimers being less abundant in full-length proteins than in the context of isolated MH2 domains. Remarkably, the presence of dimers—and not only trimers—offers a plausible explanation for the heterotrimeric association of Smad proteins in cells. In this scenario, a Smad4 (monomer) and a dimer of Smad2 can yield a Smad4-Smad2 hetero-trimer (Fig. 8). The Smad2 dimer can also associate with a monomeric Smad2 (or Smad3) to form R-Smad trimers, as observed in experiments in cells *(17–19)*. In both cases, the formation of heterotrimers starting from a dimer intermediate seems to be more favorable than when based on homo-trimer dissociation and competitive displacement triggered by Smad4, as was initially thought. Although experimental proof is needed, it is tempting to speculate that a similar mechanism of hetero-trimer formation also holds true for Smad1/5/8 proteins, whose MH1 domains are prone to define dimers, perhaps enhancing the dimerization propensity of the entire protein *(28)*.

**Fig. 8.**
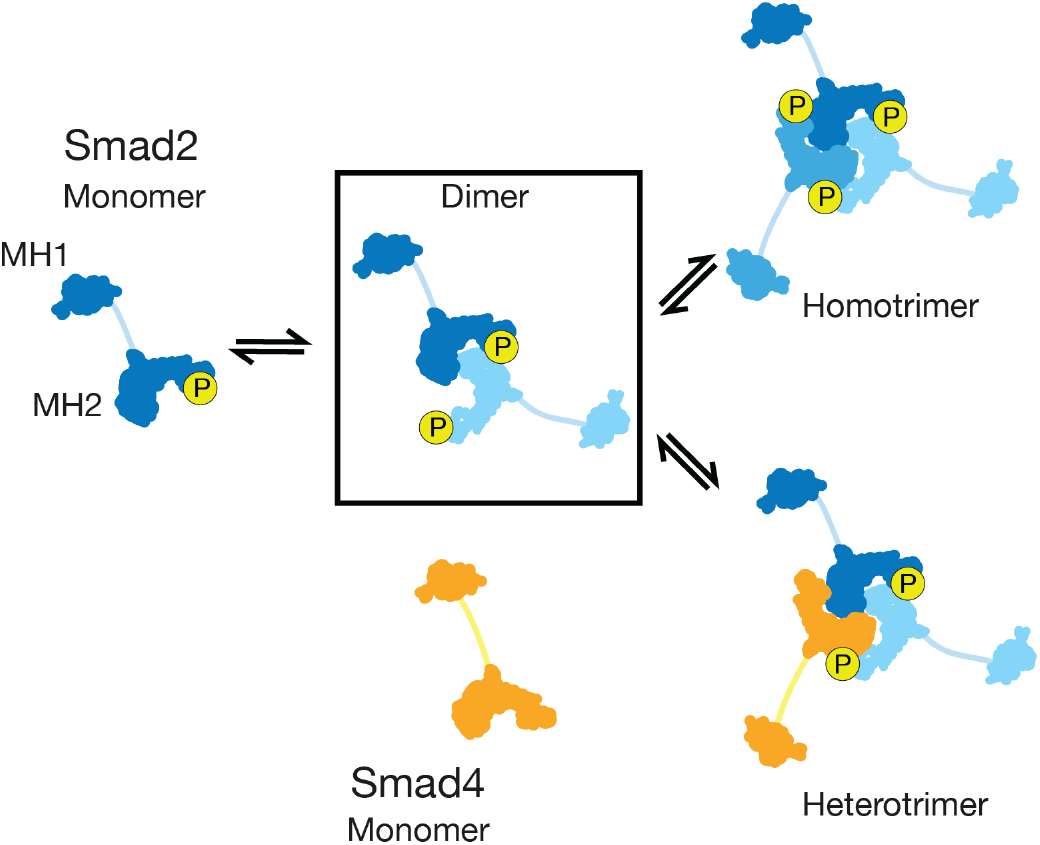
Proposed mechanism describing the hetero-trimer association of Smad proteins Schematic representation of the Smad2 and Smad4 proteins and quaternary structures. The domains are represented as silhouettes generated from 3D structures of MH1 and MH2 domains. Flexible connectors have been simplified and sketched as lines. The Smad2 dimer, (each monomer colored with a shade of blue), can associate either with monomeric Smad4, (shown in orange), and form a hetero-trimer or with another monomeric R-Smad to define homo-trimeric R-Smad assemblies.

Remarkably, in the case of Smad2FL (WT and phosphomimetic variant), we also observed an increase in compact conformations upon dimer-trimer formation. From a biological perspective, shorter inter-domain distances could act as a protective mechanism to prevent Smad2 from interacting non-specifically with other proteins, and instead, favoring oligomerization in a concentration-dependent manner. In fact, spatial and temporal variations of protein concentration in cells regulate protein function in other transcription factors, both in vivo and in vitro *(62–65)*. These results suggest that local protein concentration could act as a regulator of Smad2 activation, by modulating inter-domain distances and the transition from monomer to dimers/trimers and vice-versa. The conformational fluctuations of the linkers also limit the overall freedom of the MH1 domains by tethering them to the trimer, speeding up the process of identifying optimal binding sites through an adapted “Monkey Bar” mechanism *(66)* to sample clusters of DNA regions in *cis*-regulatory elements *(26, 28)*.

Overall, these results helped us draw the first map of the conformational landscape of full-length Smad proteins and they support the hypothesis that conformational changes in Smad proteins underlie many processes during TGFβ signaling, including the interaction of these complexes with cis-regulatory elements. These results pave the way to study the effect of modifications (such as ubiquitination and phosphorylation or disease-associated mutations) on conformational and assembly dynamics outside conserved domains. These results also help to explain domain-interactions with cofactors and with DNA in the context of full-length proteins.

Finally, we also anticipate that the methodology and computational tools optimized here will find broad application in the study of other multi-domain proteins connected by long disordered linkers, a protein class that has received little attention in structural biology due to methodological limitations.

## Materials and Methods

The experimental workflow is indicated in Fig.1A,B and explained at the beginning of the Results section.

### Recombinant protein production and cloning

S2FL, S2_460*_, S2LMH2, S2FLEEE, and S2LMH2EEE constructs were cloned in the pETM10 vector with an N-terminal His-tag, whereas S2L, S2MH1-E3, S4LMH2, S4SADMH2, S4MH2, and S4L were cloned in the pETM11 vector. S4FL was cloned in the pCOOFY34 vector with an N-terminal streptavidin tag and a 3C protease cleavage site. Successful cloning was confirmed by DNA sequencing (GATC Biotech). All constructs are shown in Fig. S1A.

Cloning was performed using standard protocols as described *(26)*. For protein production, all protein constructs were expressed in the *E. coli* BL21 (DE3) strain. Cells were cultured at 37°C in Luria-Bertani (LB) medium until reaching an OD_600_ of 0.6-0.8. After induction with IPTG (Isopropyl β-D-1-thiogalactopyranoside) at a final concentration of 0.5 mM, and overnight expression at 20°C, bacterial cultures were centrifuged at 4000x*g* for 20 min and resuspended in lysis buffer (40 mM Tris, pH 7.5, 400 mM NaCl, 0.1% Tween-20, 40 mM imidazole, 1 mM TCEP (tris(2-carboxyethyl) phosphine) and a protease inhibitor cocktail (S8820 SIGMA*FAST*™). Cells were lysed using a refrigerated EmulsiFlex-C5 (Avestin) at 20000 psi and the lysed solution was centrifuged at 35000x*g* for 45 min at 4°C to discard insoluble material. Soluble supernatants were purified by affinity chromatography (StepTag or HiTrap Chelating HP columns, GE Healthcare Life Science) using an NGC Quest 10 Plus Chromatography System (BIO-RAD) and a buffer gradient starting at 0% buffer A (lysis buffer) up to 100% buffer B in 15 column volumes. Buffer B was 40 mM Tris, pH 7.5, 200 mM NaCl, 2.5 mM desthiobiotin for S4FL and 40 mM Tris, pH 7.5, 400 mM NaCl, 400 mM imidazole for the rest. Eluted proteins were digested at 4°C with specific proteases and further purified by ion-exchange chromatography using a HiTrap SP HP or monoQ (GE Healthcare) columns and a gradient running from 0% buffer A (40 mM TRIS, pH 7.2) to 100% buffer B (40 mM TRIS, 1M NaCl, pH 7.2). As a final purification, step size-exclusion chromatography was performed using 40 mM TRIS, 150 mM NaCl, pH 7.2 buffer.

For the purification of the inter-domain linkers S2LS and S4LS, the proteins were expressed as described above but the resulting proteins were insoluble. In these cases, the lysis and protein elution were performed in denaturing conditions (40 mM TRIS, 400 mM NaCl, 8 M Urea, 40 mM imidazole, 1 mM TCEP, 0.1% tween20, pH 7.5). Proteins were refolded bound to the resin, using four washing steps and increasing the ratio of refolding/lysis buffers from zero to four (refolding buffer: 40 mM TRIS, 400 mM NaCl, 40 mM imidazole, 1 mM TCEP, 0.1% tween20, pH 7.5). After elution, proteins were cleaved and further purified by gel filtration chromatography and stored in 40 mM TRIS, 150 mM NaCl, pH 7.2. ^15^N-and ^13^C-labeled proteins were prepared as previously described *(27, 28)* and purified as above. Aliquots were kept frozen at −80 °C. Protein purity was assessed by SDS-PAGE (sodium dodecyl sulfate–polyacrylamide gel electrophoresis) and mass spectrometry.

### Nuclear magnetic resonance spectroscopy

NMR data were acquired on a Bruker Avance III 600-MHz spectrometer (IRB Barcelona) or Bruker Avance IIIHD 850-MHz (IBS-ISBG, Grenoble), both equipped with a Cryo TCI (1H, ^13^C, ^15^N, ^2^H), 5 mm, with z-gradients. S4L and S2L samples were studied in 40 mM TRIS, 150 mM NaCl, 10% D2O, pH 6.6 at 278K. The HSQC experiments were processed using TOPSPIN v3.5 (Bruker). All other experiments were processed using NMRPIPE *(67)* and analyzed with the CcpNmr Analysis *(68)* software suite or with CARA. The NMR backbone assignment followed established protocols employing CBCANH, CBCA(CO)NH, HN(COCA)NH, HN(CA)NH, HN(CA)CO, HNCO and N HCACB and NHCACOCB experiments for deuterated proteins using Non-Uniform Sampling (NUS) and BEST-TROSY backbone experiments *(69–73)*. Proline residues were connected using a set of specific experiments *(74)*. T1 and T2 relaxation measurements were acquired using standard pulse sequences at 278K *(69)*, essentially as described in *(28)*. T1 relaxation experiments used inversion recovery delays of 20, 110, 160, 270, 430, 540, 700, 860, 1080, 1400, 1720 and 2000 ms. The delays used for the T2 experiment were 0, 20, 40, 60, 80, 120, 160, 200, 280 and 400 ms. The size of the fid for all experiments was (^1^H)1024 × (^15^N)256 points and the interscan delay was set to 3s. Relaxation rates were retrieved by fitting peak intensities to an exponential function implemented in CcpNnmr analysis *(68)*.

The NMR titration of the S4MH2 into the S4MH1 construct was performed in the same buffer described above, with 1 equivalent corresponding to 130 μM.

Chemical shifts were quantified using equation 1,

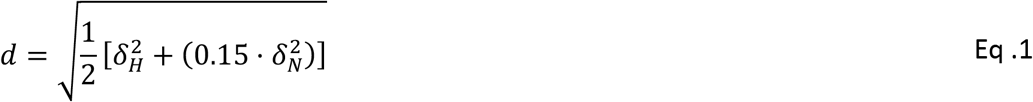

where δ_H_ and δ_N_ are the ^1^H and ^15^N chemical shift differences, respectively.

Secondary structure propensities were calculated using the ncSPC (Neighbor Corrected Structural Propensity Calculator) method *(50)* using ^13^C, ^15^N and ^1^HN backbone chemical shifts. Values between −0.1 and 0.1 are random coils, with values above 0.1 and below −0.1 corresponding to ɑ-helix and β-sheet propensity, respectively.

### Protein disorder propensities

The protein disorder propensity was calculated with MetaDisorder *(75)*. Uversky plot was calculated using CIDER *(76)*.

### Small-angle X-ray scattering data acquisition

SAXS data were acquired on Beamline 29 (BM29) at the European Synchrotron Radiation Facility (ESRF, Grenoble, France). Protein samples were centrifuged for 10 minutes at 10000x*g* prior to data acquisition. Experiments on BM29 were collected on 45 μL samples with the following settings: 12.5 keV, 100% transmission, low viscosity and 0s wait time. Data were recorded on a Pilatus 1M detector, at 10 °C. Ten frames per sample were collected for 1s each. Solvent from each sample elution was collected and their scattering data were acquired to account for buffer contribution. Image conversion to the 1D profile, scaling, buffer subtraction and radiation damage accession was done using the in-house software pipeline available at BM29. Further processing was done by the ATSAS software suite *(77)* and the ScÅtter package (http://www.bioisis.net/). For Smad4 constructs, the reported SAXS profiles were concentration-independent and were merged in ATSAS2.8 and used for subsequent analysis. For Smad2, the S2L construct was merged at the reported concentrations, and all other constructs were analyzed for each concentration individually, due to their concentration-dependence (Table 1).

**Table 1 –.**
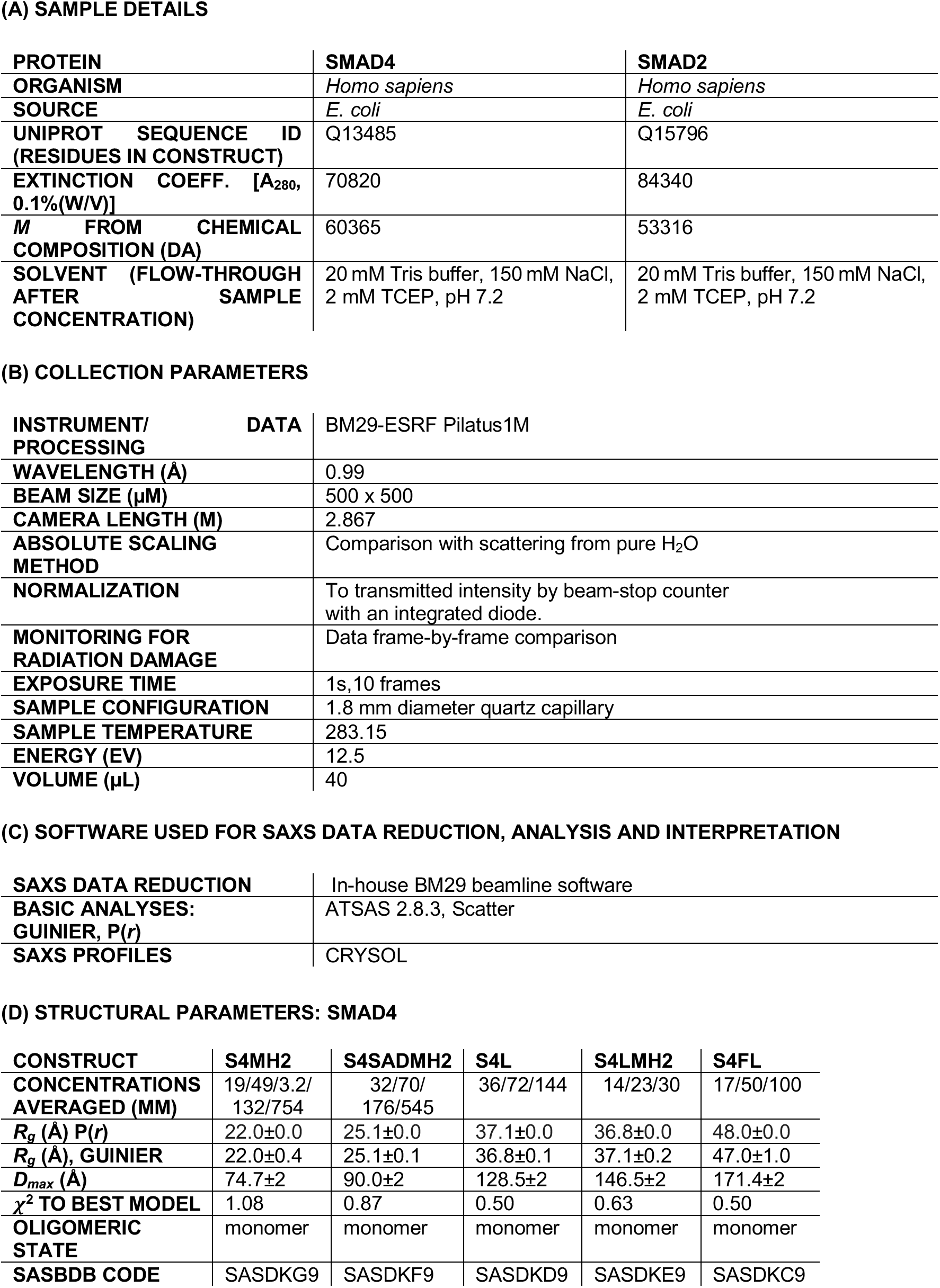
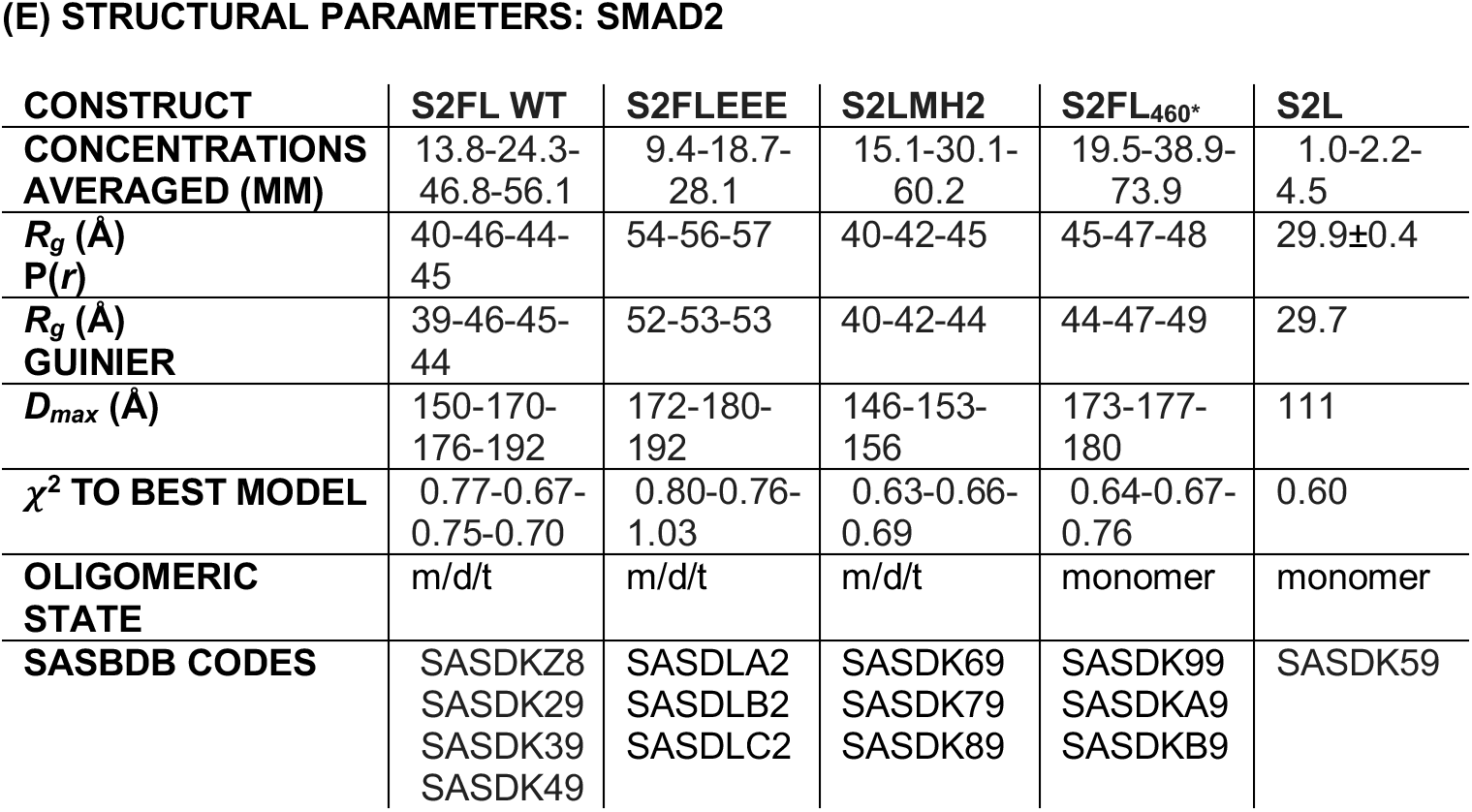
SAXS data and acquisition parameters.

### Ion mobility mass spectrometry data acquisition

Ion mobility mass spectrometry experiments were performed using a Synapt G1-HDMS mass spectrometer (Waters, Manchester, UK). Samples were buffer-exchanged to a 200 mM ammonium acetate buffer and infused by an automated chip-based nanoelectrospray using a Triversa Nanomate system (Advion BioSciences, Ithaca, NY, USA). The ionization was performed in positive mode using a spray voltage and a gas pressure of 1.70 kV and 0.5 psi, respectively. Cone voltage, extraction cone and source temperature were set to 40 V, 2 V and 20 °C, respectively. Trap and transfer collision energies were set to 10 V and 10 V, respectively. The pressure in the trap and transfer T-Wave regions were 5.8410^−2^ mbar of Ar and the pressure in the IMS T-Wave was 0.460 mbar of N_2_. Trap gas and IMS gas flows were 8 and 24 mL/sec, respectively. The travelling wave used in the IMS T-Wave for mobility separation was operated at 300 m/sec. The wave amplitude was fixed to 10 V. The bias voltage for entering in the T-wave cell was 15 V. The instrument was calibrated over the m/z range 500-8000 Da using a solution of cesium iodide. MassLynx (v4.1) and Driftscope (v2.4) were used for data processing and analysis. Drift time calibration of the T-Wave cell was performed using the following calibrants: β-Lactoglobulin (bovine milk), transthyretin (human plasma), avidin (egg white), serum albumin (bovine) and concanavalin A (*Canavalia ensiformis*) in 200 mM ammonium acetate at 20 μM. All measurements followed the same experimental protocol stated above. The reduced cross-sections (Ω’) were retrieved from previous results *(78)* and plotted against corrected drift times (tD). A power law fit of Ω’ vs. tD was performed to extract the calibration coefficients (Prism v6, GraphPad Software Inc.). Finally, Gaussian curves were fitted to the drift time distributions used to extract the experimental CCS.

### Structural modeling of Smad2 and Smad4 linkers

Random coil ensemble models of S2L and S4L containing 10,000 conformations each *(52)* were generated using *Flexible-Meccano* (FM) *(40)*, where torsion angle pairs were selected randomly from a database of amino acid-specific conformations in loop regions of high-resolution X-ray structures. Side-chains modeling with SCCOMP *(79)*, and energy-minimization in explicit solvent using GROMACS 5.1.1 were then carried out *(80)*. We used the force field AMBER99sb-ILDN *(81)* and the TIP3P water model *(82)*. We used CRYSOL *(83)* to compute the theoretical SAXS profiles from conformational ensembles of S2L and S4L. All theoretical curves were obtained with 101 points and a maximum scattering vector of 0.5 Å^-1^ using 25 harmonics. Using the ensemble optimization method (EOM) *(41)*, we select from the S2L and S4L structural pools the linker structures whose theoretical SAXS profiles collectively fit their experimental SAXS profiles, using the reduced χ^2^ metric. The theoretical SAXS profile for each generated conformation was computed and then averaged over the selected sub-ensembles.

### Reconstruction of the missing fragments of Smad2 and Smad4 MH2 domains

Ensembles of missing terminal disordered fragments were re-built using FM and attached to the X-ray templates of Smads MH1 and MH2 using *in-house* scripts as in *(84, 85)*. We used the following PDB structures as templates: Smad4 MH1 (PDB:3QSV) and MH2 (PDB:1DD1) domains, and Smad2 MH1 (PDB:6H3R) and MH2 (PDB:1KHX) domains. For each built segment, side-chains were added using SCCOMP and then pre-processed with Rosetta 3.5 *fixbb*-module to alleviate steric clashes. Internal segments and disordered loops were built using the RosettaCM application as previously described for S2MH1*(27)*, outputting 5,000 structures per domain. We assessed the quality of the ensembles by examining their averaged SAXS curves against the respective experimental data, using the reduced χ^2^ metric. We subsequently used these conformers to build LS-MH2, full-length (i.e., MH1-LS-MH2), and oligomeric constructs. All models were energy-minimized in water as above.

### Modeling and fittings of full-length Smad proteins and mutants

We modeled the full-length proteins (i.e, S4FL, S2FLWT, S2FLEEE and S2FL_460*_ variants) using the pools created for each region (i.e., MH1, MH2, MH2_460*_, and LS), explained above. To create S2FLEEE, we *in-silico* mutated the phospho-serine sites (pSer) of MH2 located at the C-terminal residues Cys-Ser-Ser-Met-Ser (SSXSS motif) by phosphomimetic glutamic acid residues. To generate ensemble models for the C-terminally-truncated Smad2FL variant (S2FL_460*_), we removed the seven last residues from the MH2 crystal structure. Different conformers were randomly selected and added to new unique explicit models without steric clashes. The final models were energy-minimized with GROMACS 5.1.1 *(80)*.

Following this workflow, we created an ensemble of 10,000 unique combinations for each system/scenario, including different oligomeric forms, i.e., their monomeric, dimer, and trimer representations. S4 and S2 MH2 domains form crystallographic homo-trimers. We used the trimers as starting models to generate monomeric and dimeric versions by removing one or two chains. Then, to probe the oligomeric preferences of Smad4 and Smad2 FL and variants, identical monomer, dimer, and trimer populations were defined in the initial shared pool and analyzed with EOM. To this end, we used CRYSOL *(83)* to compute the theoretical SAXS profiles from each conformational ensemble, and with EOM, we selected those structures reproducing the experimental SAXS data. A minimal sub-ensemble size (N_se_) was empirically determined by searching for the smallest N_se_=50 with the global lowest SAXS discrepancy (reduced χ^2^), checking for over-fitting biases. To further detail the Smad4 conformational landscape and add robustness to the modeling, we also used multiple SAXS curves from the deletion mutant S4LMH2, S4L, and full-length protein as a strategy to enhance the structural content of SAXS data and improve model discrimination. The ensemble multi-curve fitting with a single pool successfully improved the structural analysis of disordered tau protein *(86)*. Subsequent analysis of the ensembles obtained was done using MDanalysis *(87)*.

### Molecular dynamics simulations

To assess the stability of MH2 dimers, we ran a molecular dynamics trajectory of 500 ns. Molecular dynamics simulations were performed for the dimer with GROMACS 5.1.1 using the Amber99sb force field *(80)*. The system was solvated in a dodecahedron box with TIP3P water. It was minimized for a maximum of 50,000 steps or until the force constant was less than 1000 kJ/mol/nm, using the steepest descent algorithm implemented in GROMACS. The cutoff distance used for the non-bonded interactions, using the Particle mesh Ewald (PME) method, was 10 Å. Before the final production simulation, the system was equilibrated using the NPT ensemble for 500 ps, followed by 50 ps in the NVT ensemble. Finally, the system was simulated for 0.5 μs with a 2-fs integration step. The first 100 ns were discarded assuming system equilibration. Temperature coupling was done with the Nose– Hoover algorithm at 300 K. Pressure coupling was done with the Parrinello–Rahman algorithm at 1 bar. Root-mean-squared deviation (RMSD) was calculated using built-in GROMACS analysis routines and plotted using Xmgrace.

## Acknowledgments

We thank the ESRF group for help and access to the synchrotron Bio-SAXS BM29 beamline, and Dr. P. Bernardó for support and suggestions on the methodology. We thank Drs. M. Díaz and M. Vilaseca (Mass Spectrometry Core Facility, IRB Barcelona) for support with the IM-MS data and Dr. B. Brutscher (Institut de Biologie Structurale, Biomolecular NMR Spectroscopy Group, Grenoble, France) for help with the acquisition of the NMR data at 850 MHz. Dr. N. Berrow (IRB Barcelona, Protein Expression Core Facility) for his assistance with cloning and reagents, and A. Vea and C. Torner for help with protein purification. Thanks also go to Dr. J. Massagué for insightful discussions.

## Funding

This work has been financed through the Spanish MINECO program (BFU2014-53787-P and BFU2017-82675-P, M.J.M), the IRB Barcelona, the FCT – Fundação para a Ciência e a Tecnologia, I.P., Project MOSTMICRO-ITQB with references UIDB/04612/2020 and UIDP/04612/2020 (to ITQB-NOVA), FEDER Funds through COMPETE 2020 (0145-FEDER-007660), The BBVA Foundation, the Horizon 2020 Programme and the *i*NEXT program (grant Smad4 MH2 domain PID:7192).

M.J.M is an ICREA Programme Investigator and TNC is a recipient of Stimulus of Scientific Employment, Individual Support from Fundação para a Ciência e a Tecnologia, Ministério da Ciência, Tecnologia e Ensino Superior, CEECIND/01443/2017.

Access to Bio-SAXS BM29 ESRF was part of the MX-1941 BAG proposal, access to ALBA synchrotron was through the BAG proposal 2018092972.

We gratefully acknowledge institutional funding from the CERCA Programme of the Catalan Government and from the Spanish Ministry of Economy, Industry and Competitiveness (MINECO) through the Centres of Excellence Severo Ochoa award.

## Author contributions

Conceptualization: MJM, TNC

Methodology: TNC, TG, PMM

Investigation: TG, TNC, LR, EA, MJM

Supervision: MJM, TNC

Writing—review & editing: TG, PMM, TNC and MJM

## Data and materials availability

NMR data are available at the BMRB with accession codes 50738 (Smad2) and 50737 (Smad4). SAXS data are available at SASBDB, accession numbers are detailed in Table 1.

All remaining data are available in the main text or the supplementary materials.

---

**Supplementary Fig 1.**
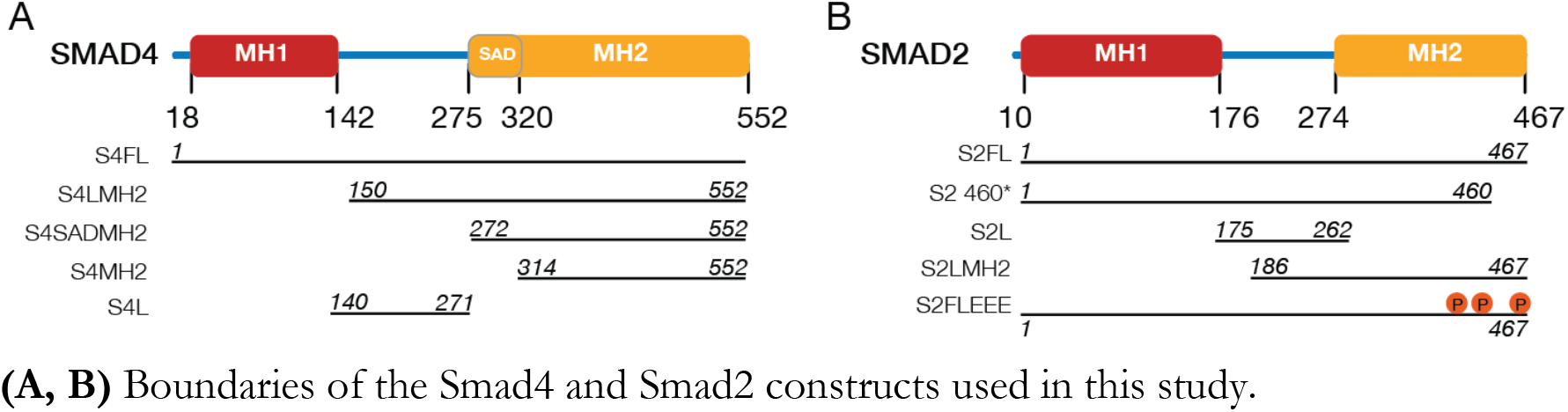
Constructs of Smad4 and Smad2 used in this study **(A, B)** Boundaries of the Smad4 and Smad2 constructs used in this study.

**Supplementary Fig 2.**
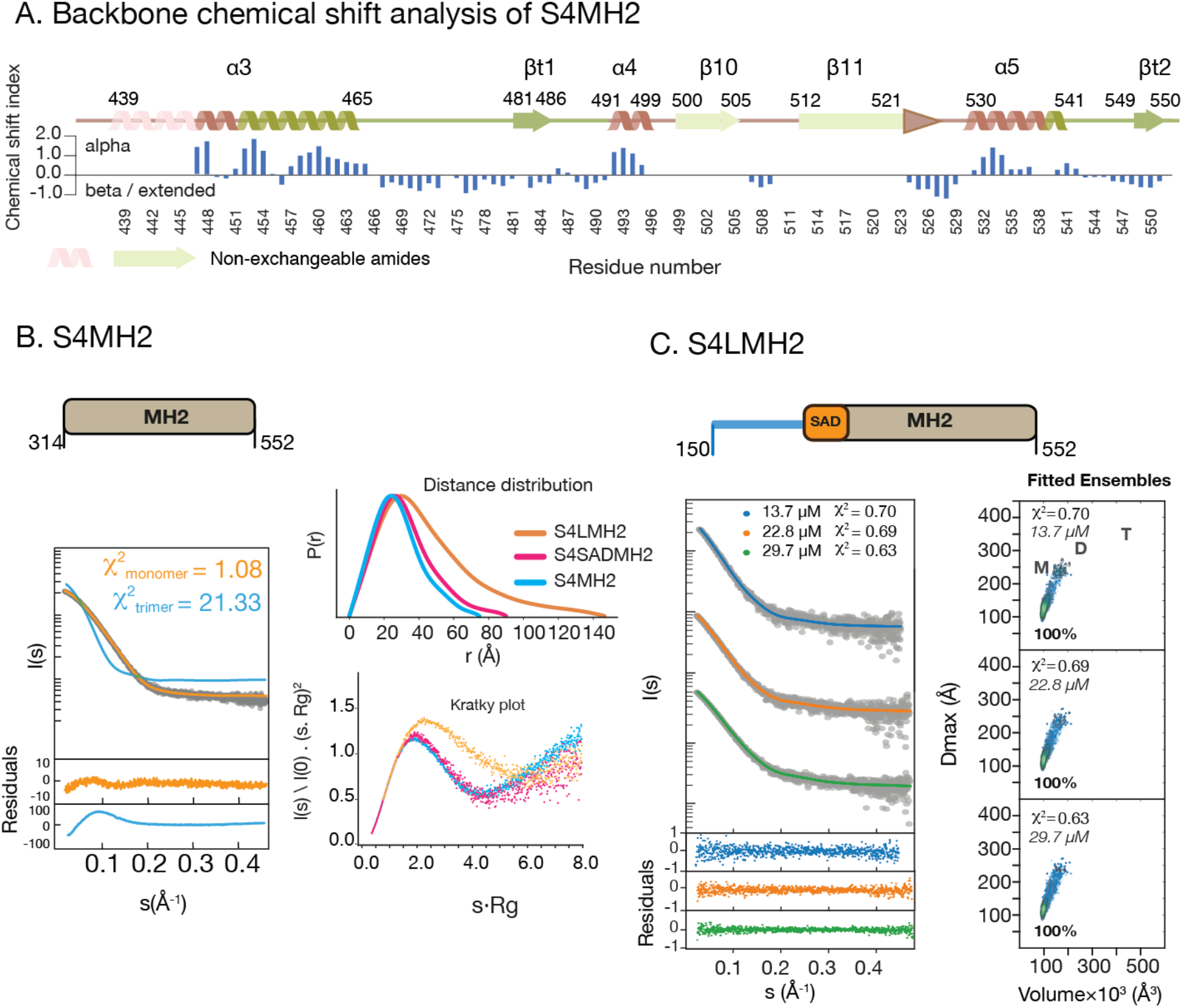
SAXS curves and fitting to EOM-derived models **(A)** Solvent-accessible areas located in and around helix3 and helix4 and in the C-terminal sequence of the MH2 domain were identified using D–H exchange and triple resonance backbone experiments. Amides located in flexible areas have a fast D–H exchange rate, thus facilitating the identification of these residues. These areas include the second half of helix3, the loop connecting to helix4 and in the beta t1 and t2 hairpin. Secondary structure propensities are represented as CS differences (positive differences indicate helical propensities whereas negative values correspond to beta or extended conformations. Secondary structure elements are colored as in Fig. 2A. **(B)** Left: SAXS curves for S4MH2 and fitting to monomer or trimer models. Residuals and χ^2^ values indicate that ensembles containing monomers but not trimers fit the experimental data. Right: Distance distributions and Kratky plot (dimensionless) derived from the SAXS experimental profiles corresponding to S4LMH2, S4SADMH2 and S4MH2. The sharp maximum observed in the Kratky plot for S4MH2 and S4SADMH2 show a globular fold of the domain. For S4LMH2, both the distance distribution asymmetry and the non-apparent maximum for the Kratky plot suggest a protein with mixed globular/disordered parts. **(C)** Left: SAXS curves for S4LMH2 at three different concentrations and fitting to EOM-derived models. Right: Kernel density contour plots for *Dmax* and Volume for each model, calculated from the EOM ensembles. M, D and T abbreviate monomer dimer and trimer species, respectively.

**Supplementary Fig 3.**
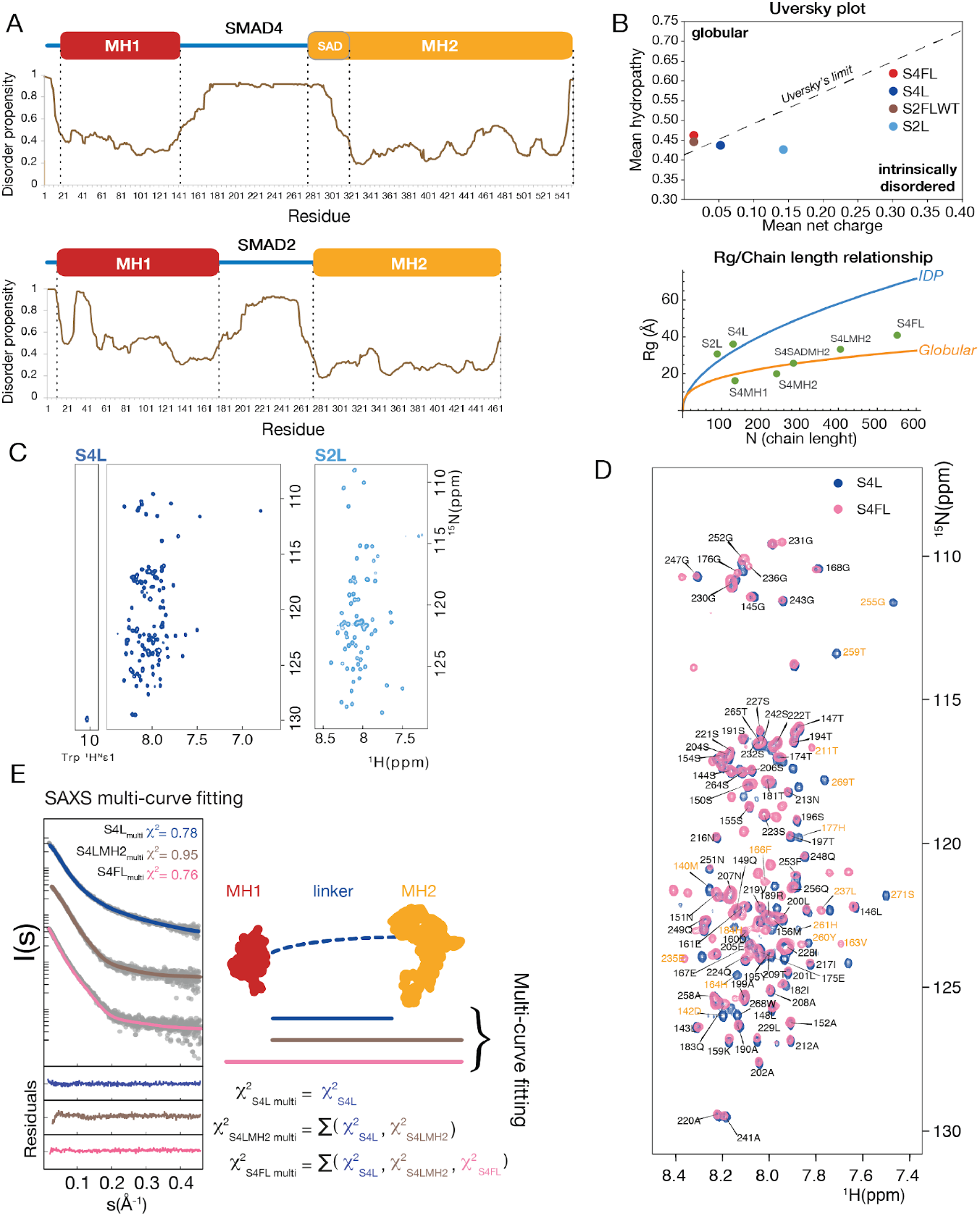
Sequence, NMR and SAXS data analysis **(A)** Disorder propensities calculated with MetaDisorder *(75)* for Smad4 and Smad2 proteins, where 0 is fully ordered and 1 fully disordered. **(B)** Uversky plot generated with CIDER *(76)* for S4FL, S4L and S2FLWT. The dashed line is the Uversky’s limit representing the partition between disordered and intrinsically disordered proteins. *Rg*/Chain length relationship for the different protein constructs used in this work. Lines correspond to Flory’s relationship parametrized for denatured proteins. Larger values than those plotted in the blue line correspond to intrinsically disordered regions *(88)*, whereas values below the orange line correspond to globular and compact systems. **(C)** ^1^H-^15^N HSQC spectra of S2L and S4L showing a narrow CS dispersion characteristic of IDRs. **(D)** Overlay of ^1^H-^15^N HSQC spectra of S4 linker in isolation or in the full-length context. No major differences are observed between the two samples. All assignments are indicated. In yellow are indicated the resonances observed for S4L but not for S4FL. **(E)** SAXS multi-curve fitting using the S4FL, S4LMH2, and S4L constructs. Conformations used for the fitting were obtained from full-length conformations by deleting the corresponding domains, and fitted to the individual SAXS curves for S4L, S4LMH2 and S4FL. Global fitting 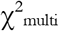 is the best-fit curve given by the EOM ensemble using the three SAXS profiles simultaneously.

**Supplementary Fig 4.**
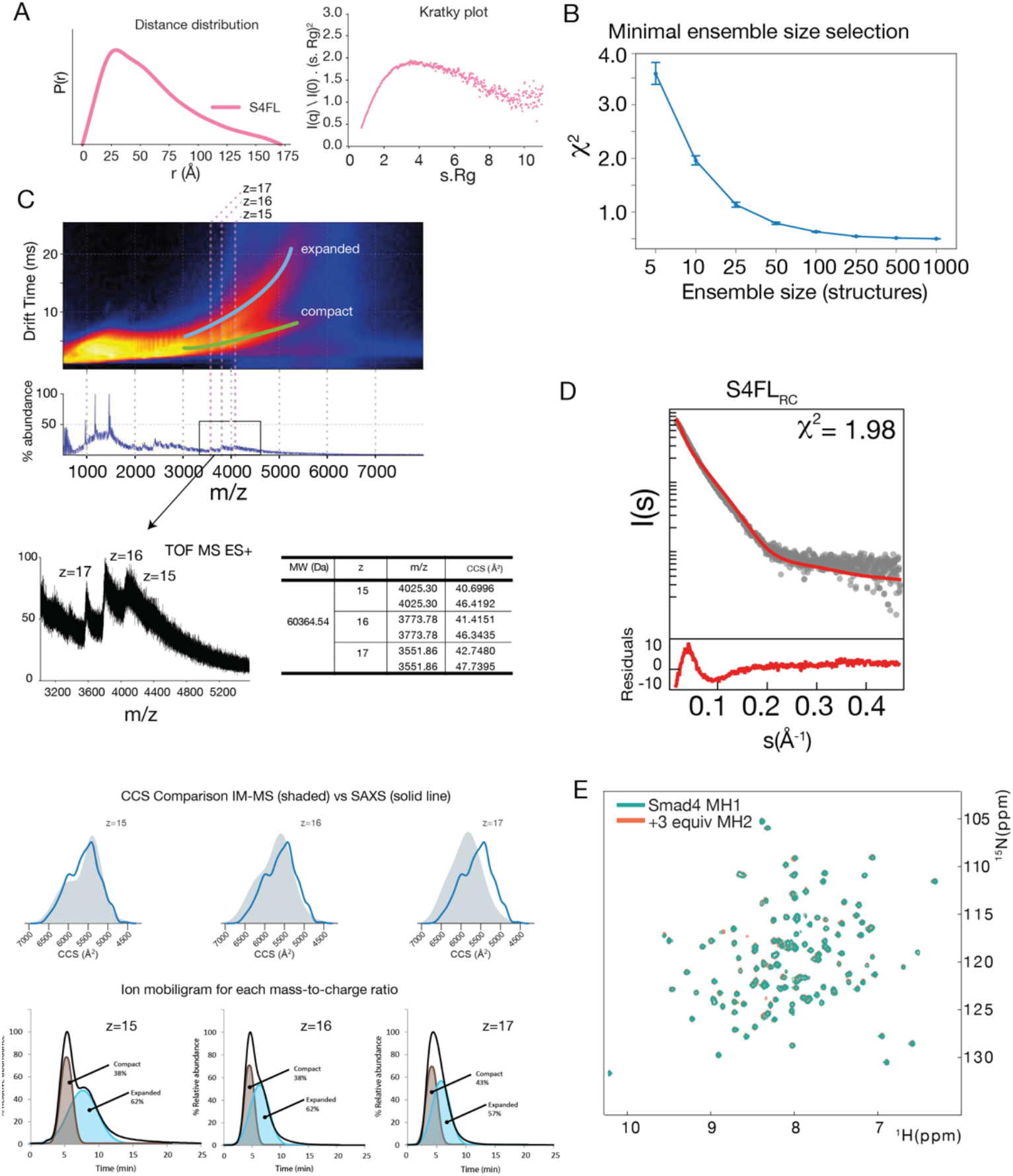
SAXS, NMR and IMMS data analysis for S4FL. **(A)** Pair distance distributions and Kratky plot showing a profile in agreement with a mixed globular/flexible protein. **(B)** Grid-search selection to determine the minimal size of ensembles that fulfill the experimental data to avoid overfitting. The graph represents sub-ensembles of different sizes and χ^2^ values. Sub-ensembles in the range of 30–50 conformers fulfilled the condition of fitting the SAXS data with a χ^2^ lower than 1.0. We selected sub-ensembles of 50 conformations in this study. **(C)** Plot of the mobility drift time versus m/z (*Top)*. Compact and expanded forms are indicated in the figure. Below, the ion mobiligrams for three selected peaks (z=15,16 and 17) are shown and the compact and expanded components are integrated, showing ratios similar to those obtained by SAXS by simulating the Collision cross-sections for the EOM ensemble, calculated with the IMPACT software *(89)* using the PA method (*Bottom*). **(D)** Negative control of the analysis: SAXS fitting (red) to the ensemble before EOM selection yielded a non-uniform residual distribution and poor fitting. **(E)** ^1^H-^15^N HSQC CS differences induced in the S4MH1 domain after the addition of the S4MH2 domain. The differences are almost negligible.

**Supplementary Fig 5.**
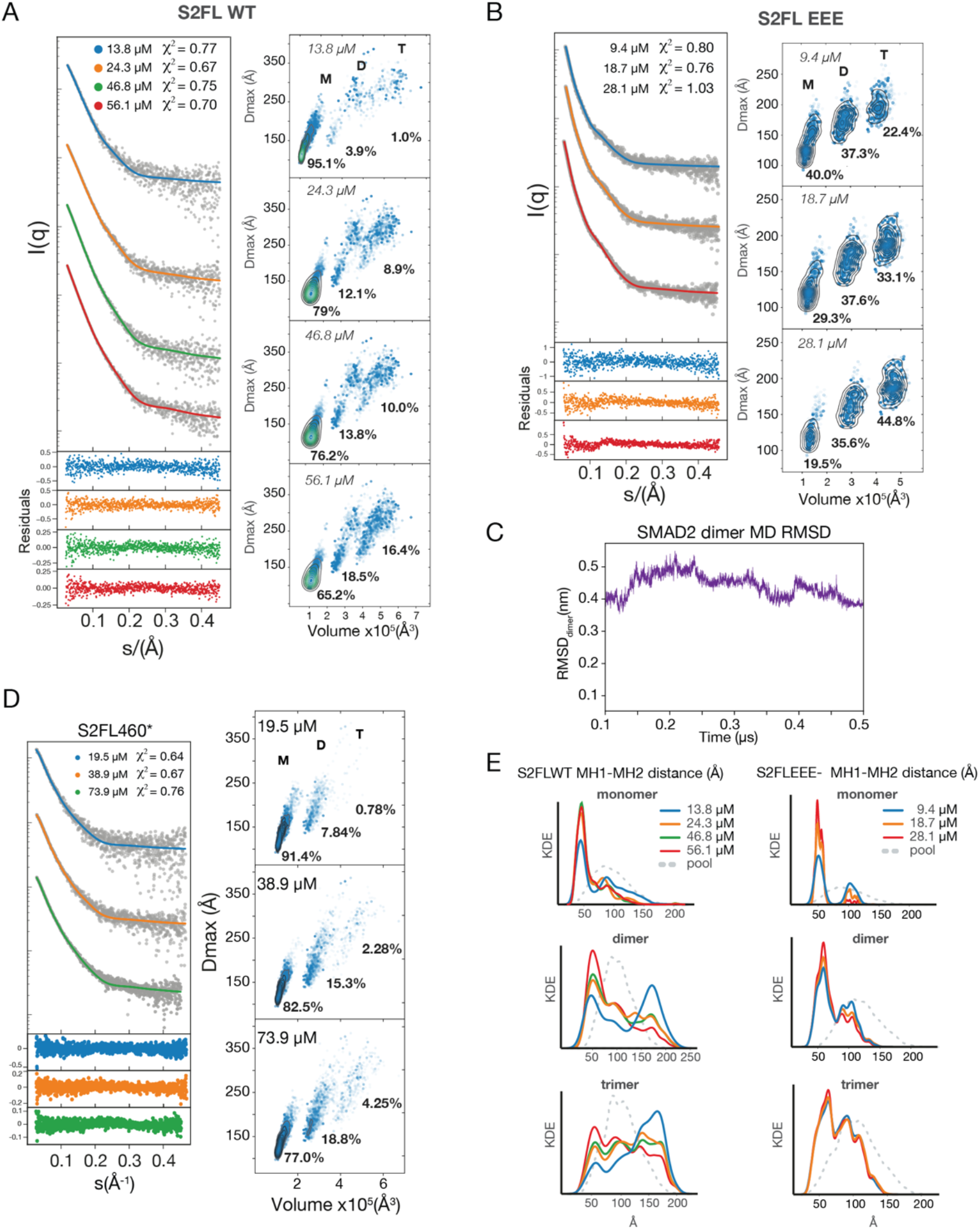
SAXS curves and inter-domain distances retrieved from the EOM ensemble. **(A** and **B)** Left: SAXS curves for S2FL WT(A) or S2FL EEE(B) at different concentrations with the fitting to EOM-derived models. Right: Kernel density contour plots for *Dmax* and Volume for each model, calculated from the EOM ensembles. M, D, and T are monomer, dimer, and trimer species, respectively. **(C)** MD simulations of S2MH2 dimer (0.5 μs). The solid line represents the Residue mean squared deviation (RMSD) of the homodimer along the trajectory. **(D)** Left: SAXS curves for S2FL460* at different concentrations and fitting to EOM-derived models. Right: Kernel density contour plots for *Dmax* and Volume for each model, calculated from the EOM ensembles. M, D, and T are monomer, dimer, and trimer species, respectively. **(E)** Inter-domain distances retrieved from the SAXS EOM ensemble for monomer, dimer, and trimer derived from panels A and B. The random coil ensemble is shown in gray dashed lines.

